# A combined *in silico*, *in vitro* and clinical approach to characterise novel pathogenic missense variants in PRPF31 in retinitis pigmentosa

**DOI:** 10.1101/480343

**Authors:** Gabrielle Wheway, Liliya Nazlamova, Nervine Meshad, Samantha Hunt, Nicola Jackson, Amanda Churchill

## Abstract

At least six different proteins of the spliceosome, including PRPF3, PRPF4, PRPF6, PRPF8, PRPF31 and SNRNP200, are mutated in autosomal dominant retinitis pigmentosa (adRP). These proteins have recently been shown to localise to the base of the connecting cilium of the retinal photoreceptor cells, elucidating this form of RP as a retinal ciliopathy. In the case of loss-of-function variants in these genes, pathogenicity can easily be ascribed. In the case of missense variants, this is more challenging. Furthermore, the exact molecular mechanism of disease in this form of RP remains poorly understood.

In this paper we take advantage of the recently published cryo EM-resolved structure of the entire human spliceosome, to predict the effect of a novel missense variant in one component of the spliceosome; PRPF31, found in a patient attending the genetics eye clinic at Bristol Eye Hospital. Monoallelic variants in *PRPF31* are a common cause of autosomal dominant retinitis pigmentosa (adRP) with incomplete penetrance. We use *in vitro* studies to confirm pathogenicity of this novel variant *PRPF31* c.341T>A, p.Ile114Asn.

This work demonstrates how *in silico* modelling of structural effects of missense variants on cryo-EM resolved protein complexes can contribute to predicting pathogenicity of novel variants, in combination with *in vitro* and clinical studies. It is currently a considerable challenge to assign pathogenic status to missense variants in these proteins.

## 1 Introduction

Retinitis pigmentosa (RP) is a progressive retinal degeneration characterised by night blindness and restriction of peripheral vision. Later in the course of the disease, central and colour vision can be lost. Many patients experience the first signs of RP between 20-40 years but there is much phenotypic variability from age of onset and speed of deterioration to severity of visual impairment (Hartong *et al.*, 2006).

RP, whilst classified as a rare disease, is the most common cause of inherited blindness worldwide. It affects between 1:3500 and 1:2000 people (Golovleva *et al.*, 2010; Sharon and Banin, 2015), and can be inherited in an autosomal dominant (adRP), autosomal recessive (arRP), or X-linked (xlRP) manner. It may occur in isolation (non-syndromic RP) (Verbakel *et al.*, 2018), or with other features (syndromic RP) as in Bardet-Biedl syndrome, Joubert syndrome and Usher syndrome (Mockel *et al.*, 2011).

The condition is extremely heterogeneous, with 64 genes identified as causes of non-syndromic RP, and more than 50 genes associated with syndromic RP (RetNet https://sph.uth.edu/retnet/sum-dis.htm). Even with current genetic knowledge, diagnostic detection rate in adRP cohorts remains between 40% (Mockel *et al.*, 2011) and 66% (Zhang *et al.*, 2016), suggesting that many disease genes remain to be identified, and many mutations within known genes require characterization to ascribe pathogenic status. Detection rates are as low as 14% in cohorts of simplex cases (single affected individuals) and multiplex cases (several affected individuals in one family but unclear pattern of inheritance) (Jin *et al.*, 2008). Such cases account for up to 50% of RP cases, so this presents a significant challenge to diagnosis (Greenberg *et al.*, 1993; Haim, 1993; Najera *et al.*, 1995).

The second most common genetic cause of adRP is *PRPF31*, accounting for 6% of US cases (Sullivan *et al.*, 2013) 8% of Spanish cases (Martin-Merida *et al.*, 2018), 8% of French Canadian cases (Coussa *et al.*, 2015), 8% of French cases (Audo *et al.*, 2010), 8.9% of cases in North America (Daiger *et al.*, 2014), 11.1% in small Chinese cohort (Lim *et al.*, 2009), 10% in a larger Chinese cohort (Xu *et al.*, 2012) and 10.5% of Belgian cases (Van Cauwenbergh *et al.*, 2017). However, this is likely to be an underestimate due to variable penetrance of this form of RP, complicating attempts to co-segregate the variant with clinical disease, making genetic diagnosis difficult.

Whilst the majority of reported variants in *PRPF31* are indels, splice site variants and nonsense variants, large-scale deletions or copy number variations (Martin-Merida *et al.*, 2018), which are easily ascribed pathogenic status, at least eleven missense variants in *PRPF31* have been reported in the literature (**Table 1**). Missense variants are more difficult to characterize functionally than nonsense or splicing mutations (Cooper and Shendure, 2011) and it is likely that there are false negative diagnoses in patients carrying missense mutations due to lack of confidence in prediction of pathogenicity of such variants. This is reflected in the enrichment of *PRPF31* missense variants labelled ‘uncertain significance’ in ClinVar, a public repository for clinically-relevant genetic variants (Landrum *et al.*, 2016; Landrum *et al.*, 2014). Furthermore, work has shown that some variants annotated as missense *PRPF31* variants may in fact be affecting splicing of *PRPF31*, introducing premature stop codons leading to nonsense mediated decay (NMD), a common disease mechanism in RP11 (Rio Frio *et al.*, 2008). One example is c.319C>G, which, whilst originally annotated as p.Leu107Val, actually affects splicing rather than an amino acid substitution (Rio Frio *et al.*, 2008). The presence of exonic splice enhancers is often overlooked by genetics researchers.

**Table 1.**
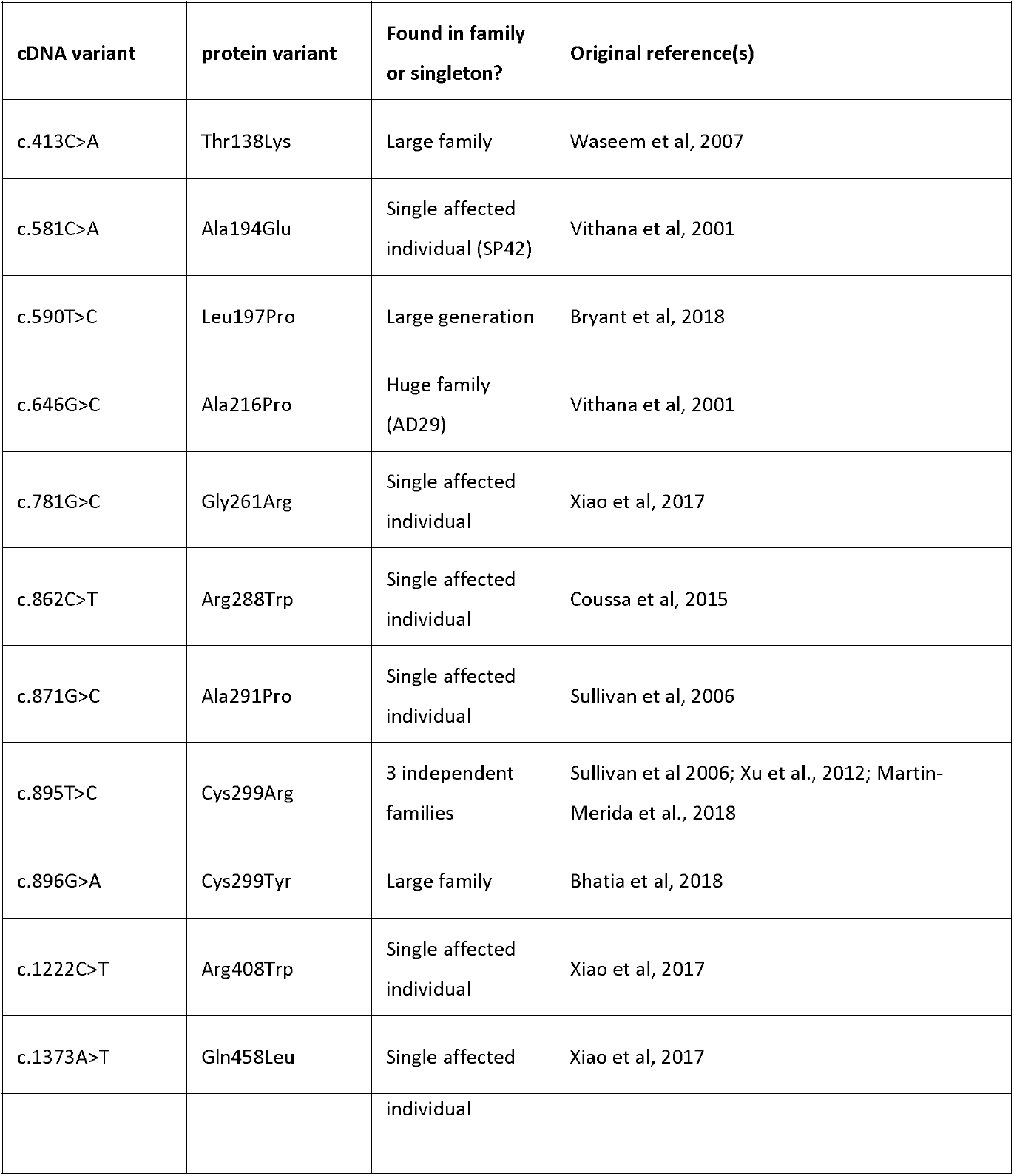
Summary of published missense mutations in *PRPF31*

| cDNA variant | protein variant | Found in family or singleton? | Original reference(s) |
| --- | --- | --- | --- |
| c.413C>A | Thr138Lys | Large family | Waseem et al, 2007 |
| c.581C>A | Ala194Glu | Single affected individual (SP42) | Vithana et al, 2001 |
| c.590T>C | Leu197Pro | Large generation | Bryant et al, 2018 |
| c.646G>C | Ala216Pro | Huge family (AD29) | Vithana et al, 2001 |
| c.781G>C | Gly261Arg | Single affected individual | Xiao et al, 2017 |
| c.862C>T | Arg288Trp | Single affected individual | Coussa et al, 2015 |
| c.871G>C | Ala291Pro | Single affected individual | Sullivan et al, 2006 |
| c.895T>C | Cys299Arg | 3 independent families | Sullivan et al 2006; Xu et al., 2012; Martin-Merida et al., 2018 |
| c.896G>A | Cys299Tyr | Large family | Bhatia et al, 2018 |
| c.1222C>T | Arg408Trp | Single affected individual | Xiao et al, 2017 |
| c.1373A>T | Gln458Leu | Single affected | Xiao et al, 2017 |

**Table 1. Summary of published missense mutations in *PRPF31***
|  |  |  |
| --- | --- | --- |
|  |  | individual |

PRPF31 is a component of the spliceosome, the huge macromolecular ribonucleoprotein (RNP) complex which catalyses the splicing of pre-messenger RNAs (pre-mRNAs) to remove introns and produce mature mRNAs (Will and Luhrmann, 2011). The spliceosome is composed of 5 small nuclear RNAs (snRNAs), U1-U5, and many proteins including pre-mRNA splicing factors PRPF3, PRPF4, PRPF6, PRPF8, and SNRNP200, all of which are also genetic causes of RP (Ruzickova and Stanek, 2016). It is unclear whether variants in these proteins have an effect on splicing of specific retinal transcripts (Deery *et al.*, 2002; Yuan *et al.*, 2005; Mordes *et al.*, 2007; Wilkie *et al.*, 2008). Some papers have failed to find any evidence for a generalized RNA splicing defects (Rivolta *et al.*, 2006). Pre-mRNA splicing factors may have additional roles beyond splicing in the nucleus, after a study recently found that PRPF6, PRPF8 and PRPF31 are all localized to the base of the retinal photoreceptor connecting cilium and are essential for ciliogenesis, suggesting that this form of RP is a ciliopathy (Wheway *et al.*, 2015). Missense variants in these proteins are, collectively, a common cause of adRP. This presents significant challenges in providing accurate diagnosis for patients with missense variants in these genes. Developing tools to provide accurate genetic diagnoses in these cases is a significant clinical priority.

The most commonly used *in silico* predictors of pathogenicity of missense variants, PolyPhen2 (Adzhubei *et al.*, 2010) and CADD (Kircher *et al.*, 2014), which use combined sequence conservation, structural and machine learning techniques only have around 15 – 20% success rate in predicting truly pathogenic variants (Miosge *et al.*, 2015). Use of simple tools has around the same success rate (Gnad *et al.*, 2013), and use of several tools in combination increases reliability (Gonzalez-Perez and Lopez-Bigas, 2011). Insight from structural biologists and molecular cell biologists is essential to make accurate predictions.

In this study we take advantage of the recently elucidated structure of the in-tact spliceosome to model the effect of a novel variant in *PRPF31,* found in a patient attending the genetics eye clinic at Bristol Eye Hospital. We combine this *in silico* analysis with *in vitro* studies to characterize this novel variant. We show that analysis of protein complexes *in silico* can complement clinical and laboratory studies in predicting pathogenicity of novel genetic variants.

## Methods

### Genetic testing

The study was conducted in accordance with the Declaration of Helsinki. Informed consent for diagnostic testing was obtained from the proband in clinic. Genomic DNA was extracted from a peripheral blood sample by Bristol Genetics Laboratory and tested against the retinal dystrophy panel of 176 genes in the NHS accredited Genomic Diagnostics Laboratory at Manchester Centre for Genomic Medicine, UK.

### Splicing analysis

We used Human Splicing Finder (Desmet *et al.*, 2009) to identify and predict the effect of variants on splicing motifs, including the acceptor and donor splice sites, branch point and auxiliary sequences known to enhance or repress splicing. This programme uses 12 different algorithms to make a comprehensive prediction of the effect of variants on splicing.

### 3D structural protein analysis

PyMol (Schrodinger Ltd) programme was used to characterize the effect of missense variants in human *PRPF31* protein. Missense variants were modelled on PRPF31 in the pre-catalytic spliceosome primed for activation (PDB file 5O9Z) (Bertram *et al.*, 2017).

### Variant construct cloning

Full-length, sequence-validated *PRPF31* ORF clone with C-terminal myc tag was obtained from Origene. c.341T>A variant was introduced using NEB Q5 site-directed mutagenesis kit. The entire wild-type and mutant clone sequence was verified by Sanger sequencing (Source Bioscience).

### Cell culture

HEK293 cells were cultured in DMEM high glucose + 10% FCS at 37°C, 5% CO_2_, and split at a ratio of 1:8 once per week. hTERT-RPE1 cells (ATCC CRL-4000) were cultured in DMEM/F12 (50:50 mix) + 10% FCS at 37°C, 5% CO_2_, and split at a ratio of 1:8 once per week.

### Cell transfection

The construct was transfected into HEK293 cells using PEI, and into hTERT-RPE1 cells using the Lonza Nucleofector.

### Inhibition of protein translation

Cells were grown for 72 hours, and treated with 30µg/ml cycloheximide in DMSO. Untreated cells were treated with the equivalent volume of DMSO.

### Protein extraction

Total protein was extracted from cells using 1% NP40 lysis buffer and scraping. Insoluble material was pelleted by centrifugation at 10,000 x g. Cell fractionation was carried out by scraping cells into fractionation buffer containing 1mM DTT, and passed through a syringe 10 times. Nuclei were pelleted at 720 x g for 5 minutes and separated from the cytoplasmic supernatant. Insoluble cytoplasmic material was pelleted using centrifugation at 10,000 x g for 5 minutes. Nuclei were washed, and lysed with 0.1% SDS and sonication. Insoluble nuclear material was pelleted using centrifugation at 10,000 x g for 5 minutes.

### SDS-PAGE and western blotting

20µg of total protein per sample with 2 x SDS loading buffer was loaded onto pre-cast 4-12% Bis-Tris gels (Life Technologies) alongside Spectra Multicolor Broad range Protein ladder (Thermo Fisher). Samples were separated by electrophoresis. Protein was transferred to PVDF membrane. Membranes were incubated with blocking solution (5% (w/v) non-fat milk/PBS), and incubated with primary antibody overnight at 4°C. After washing, membranes were incubated with secondary antibody for 1 hour at room temperature and exposed using 680nm and/or 780nm laser, or incubated with SuperSignal West Femto reagent (Pierce) and exposed using Chemiluminescence settings on LiCor Odyssey imaging system (LiCor).

### Primary antibodies for WB

Mouse anti β actin clone AC-15. 1:4000. Sigma-Aldrich A1978

Goat anti-PRPF31 primary antibody 1:1000 (AbNova)

Mouse anti-c myc 1:5000 (Sigma)

Mouse anti PCNA-HRP conjugated 1:1000 (BioRad)

### Secondary antibodies for WB

Donkey anti mouse 680 1:20,000 (LiCor)

Donkey anti goat 800 1:20,000 (LiCor)

### Immunocytochemistry

Cells were seeded at 1 × 10^5^ per mL on sterile glass coverslips in complete media. Media was changed to serum-free media after 48 hours, and cells were grown for a further 72 hours. Cells were fixed in ice-cold methanol at −20°C for 5 minutes, immediately washed with PBS, and incubated with blocking solution (1% w/v non-fat milk powder/PBS). Coverslips were incubated with primary antibodies at 4°C overnight and with secondary antibodies and DAPI for 1 hour at room temperature. Cells were mounted onto slides with Mowiol.

### Primary antibodies for IF

Goat anti-PRPF31 primary antibody 1:200 (AbNova)

Mouse anti-c myc 1:1000 (Sigma)

### Secondary antibodies for IF

Donkey anti mouse IgG AlexaFluor 488 1:500

Donkey anti goat IgG AlexaFluor 633 1:500

### Confocal imaging

Confocal images were obtained at the Centre for Research in Biosciences Imaging Facility at UWE Bristol, using a HC PL APO 63x/1.40 oil objective CS2 lens on a Leica DMi8 inverted epifluorescence microscope, attached to a Leica SP8 AOBS laser scanning confocal microscope. Images were captured using LASX software, assembled in Adobe Photoshop, and figures prepared using Adobe Illustrator.

## Results

### Clinical description of c.341T>A p.Ile114Asn patient

A 39 year old female presented to the Genetic Eye clinic at Bristol Eye Hospital in 2013 complaining of some difficulty with dark adaptation, driving at night and a reduction in her field of vision (having to turn her head to see her children). She described other family members having similar symptoms and losing their sight at a relatively young age (**Figure 1a**). Her general health was otherwise good. Over a 4 year period her best corrected visual acuity remained good at 6/6-3 right eye and 6/7.5 left eye (Snellen equivalent using a LogMar chart) whilst her peripheral vision deteriorated from an isolated mid-peripheral scotoma to tunnel vision by 2017. Fundoscopy showed widespread bilateral bone spicule pigmentation, attenuated retinal vessels and pale optic nerves typical of RP (**Figure 1b**). There was no evidence of lens opacities or macula oedema in either eye.

**Figure 1.**
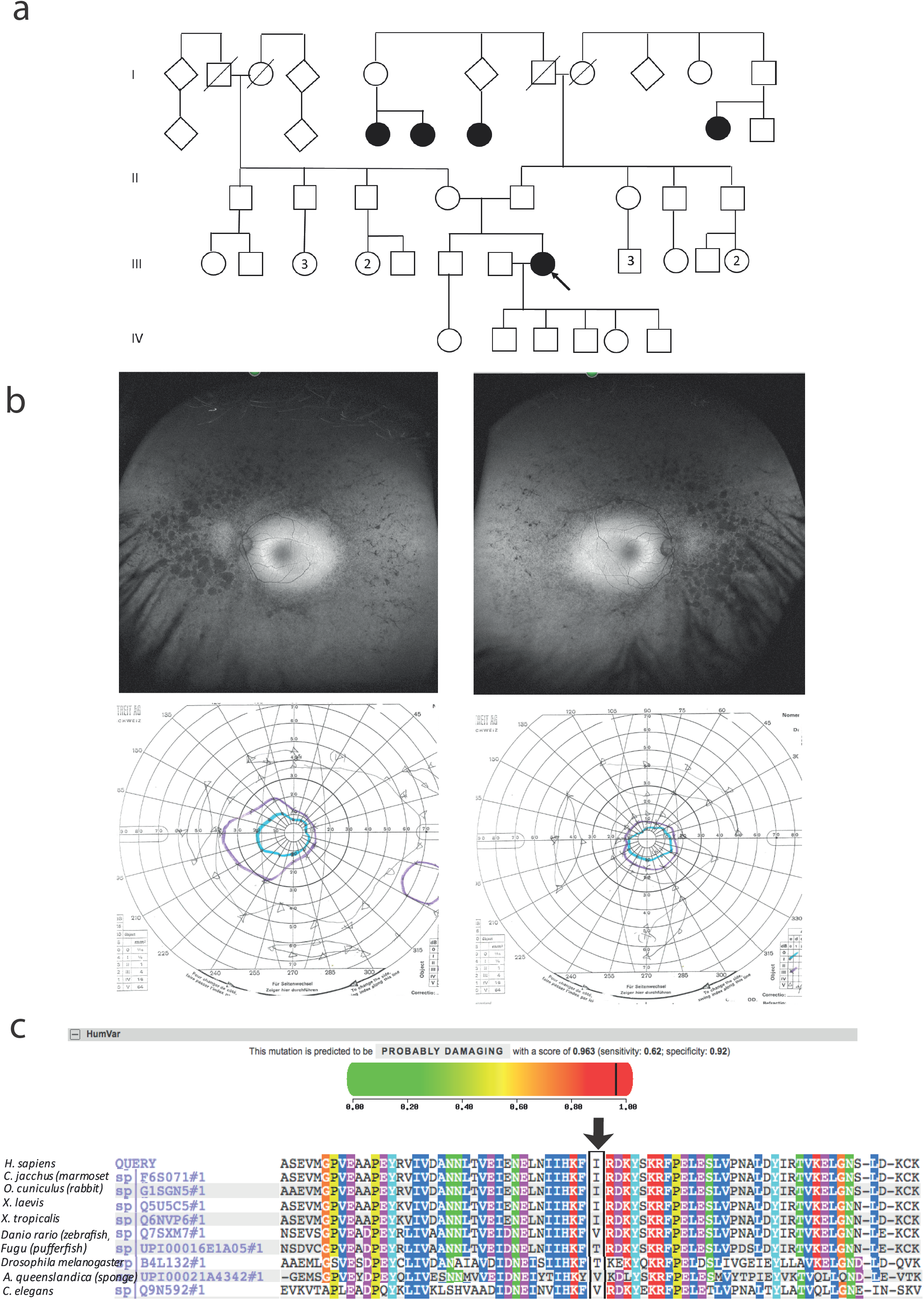
Clinical characteristics of patient with *PRPF31* c.341T>A Ile114Asn variant, and PolyPhen2 and conservation analysis of the variant **(a)** Family pedigree. Affected individuals in generation II had visual symptoms suggestive of retinitis pigmentosa and appear on both sides of the paternal grandparents of the proband. Arrow = proband **(b)** Images of clinical investigations conducted at visit in 2017. Upper panel: Red free fundus photographs show extensive bilateral retinal pigment disruption, especially nasally. Lower panel: Goldmann visual field images show bilateral tunnel vision with a small island of peripheral vision in the right eye. **(c)** PolyPhen2 score predicts this variant is probably damaging with a score of 0.963 (top), alignment of PRPF31 sequence showing conservation of Ile114 and surrounding amino acids (bottom). Ile114 identity is conserved across tetrapods, from human to *Xenopus tropicalis*, and non-polar hydrophobic similarity is conserved from yeast to human, with variations in highly derived insects (*Drosophila melanogaster*) and fish (*Fugu*).

### Variant Analysis of c.341T>A p.Ile114Asn

A heterozygous *PRPF31* change, c.341T>A p.(Ile114Asn) was identified which was confirmed by bidirectional Sanger sequencing. This variant is not present in the heterozygous or homozygous state in any individuals within the gnomAD database, nor are any other variants affecting Ile114, suggesting that this is a highly conserved residue. Analysis by PolyPhen2 suggested this change was probably damaging, with a score of 0.963 (**Figure 1c**) and SIFT concurred with this prediction with a score of 0.0. Comparative genomic alignment shows the residue to be conserved from humans to amphibia, within a highly conserved region, conserved across diverse metazoa including sponges (**Figure 1c**).

### Splicing analysis of genetic single nucleotide variants in *PRPF31*

We undertook *in silico* splicing analysis of our novel variant of interest c.341T>A p. Ile114Asn and found that it was not predicted to affect splicing. We also studied the nine published variants in *PRPF31* annotated as missense, and interestingly, five were predicted to potentially alter splicing, and one (c.1373A>T, p. Gln458Leu (Xiao *et al.*, 2017)) was predicted to be highly likely to affect splicing (**Table 2**). This suggests that either this splice predictor should be used with caution, or that p.Gln458Leu may be mis-annotated as a missense variant, when it actually affects splicing. We suggest that this variant should be a priority for further functional characterization *in vitro*.

**Table 2.** Mutations in PRPF31 annotated as missense, and their predicted impact on splicing All published missense mutations in *PRPF31*, and their predicted impact on splicing, according to Human Splicing Finder.

| cDNA variant | protein variant | Predicted effect on splicing (Human Splicing Finder) | Notes | Summary - effect on splicing? | Estimate of pathogenicity |
| --- | --- | --- | --- | --- | --- |
| c.413C>A | Thr138Lys | Potential alteration of splicing |  | Maybe | Pathogenic |
| c.581C>A | Ala194Glu | Potential alteration of splicing | Functional characterisation shows functional effect of missense change | No | Pathogenic |
| c.590T>C | Leu197Pro | Potential alteration of splicing | Functional characterisation shows functional effect of missense change | No | Pathogenic |
| c.646G>C | Ala216Pro | Potential alteration of splicing | Functional characterisation shows functional effect of missense change | No | Pathogenic |
| c.781G>C | Gly261Arg | No impact on splicing |  | No | Pathogenic |
| c.862C>T | Arg288Trp | Potential alteration of splicing |  | Maybe | Pathogenic |
| c.871G>C | Ala291Pro | No impact on splicing |  | No | Pathogenic |
| c.895T>C | Cys299Arg | No impact on splicing |  | No | Pathogenic |
| c.896G>A | Cys299Tyr | Potential alteration of splicing |  | Maybe | Pathogenic |
| c.1222C>T | Arg408Trp | Potential alteration of splicing |  | No | Pathogenic |
| c.1373A>T | Gln458Leu | **Most probably affecting splicing** |  | **Yes** | Pathogenic |

### 3D structural analysis of missense variants in PRPF31

We mapped all published missense variants onto the PRPF31 protein structure in the pre-catalytic spliceosome. For simplicity, we only show PRPF31 in complex with U4 snRNA and 15.5K (SNU13) protein (**Figure 2**) and (in complex with PRPF6 in **Supplementary Figure 1**; in complex with PRPF8 in **Supplementary Figure 2**). This showed that variants are located throughout the protein, but concentrated in several key domains. Three variants (Arg288Trp, Ala291Pro and Cys299Arg), are located in α-helix 12 of the protein, in the Nop domain which interacts with RNA and the 15.5K (SNU13) protein. Three variants are in α-helix 6 of the coiled-coil domain (Ala194Glu, Leu197Pro, Ala216Pro) and one variant is in α-helix 3 of the protein in the coiled-coil tip (Thr138Lys). Gly261Arg is within the flexible loop between the Nop and coiled-coil domains and Arg408Trp alone is in the C-terminal domain.

**Figure 2.**
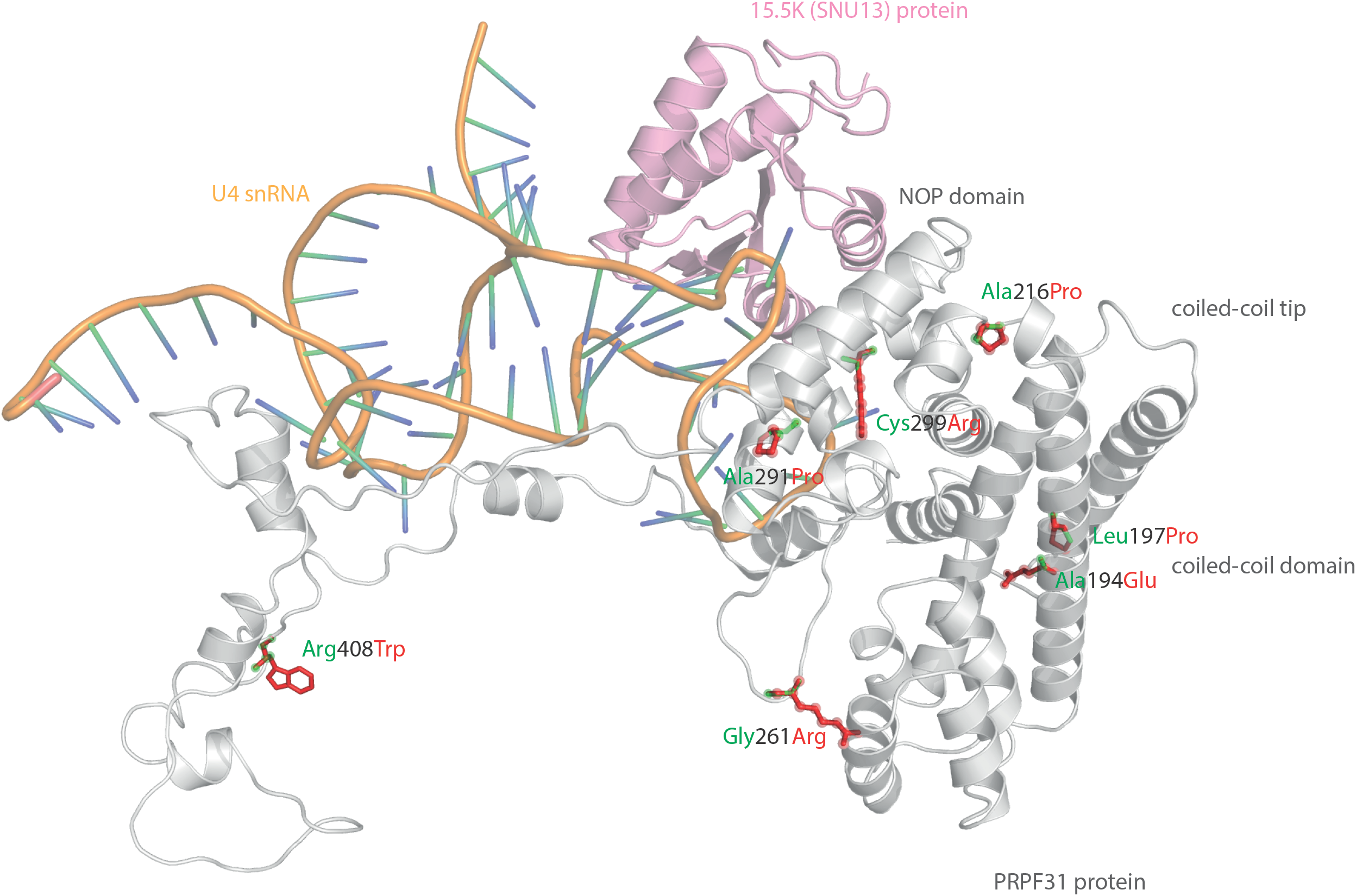
3D cartoon representation of PRPF31, including published missense mutations Cartoon representation of alpha helical structure of PRPF31 (grey) and 15.5K/SNU13 (pink) with U4 snRNA (orange backbone), with published missense mutations mapped onto the physical structure, with wild-type amino acid structure in green, and mutant amino acid structure overlaid in red.

Analysis of interactions within 4Å of each amino acid show that in most cases (Thr138Lys, Ala194Glu, Gly261Arg, Arg288Trp, Ala291Pro and Cys299Arg), these substitutions are predicted to affect hydrogen (H) bonding in PRPF3. H bonds with donor-acceptor distances of 2.2-2.5 Å are strong and mostly covalent; 2.5-3.2 Å are moderate mostly electrostatic and 3.2-4 are weak electrostatic interactions and can be predicted to be affecting protein folding and solubility (Jeffrey 1997). In the case of Arg408Trp, the substitution does not affect H bonding within PRPF31, but does introduce a new interaction with neighbouring PRPF6 (**Figure 3a; Supplementary Figures 1 and 2**). Gly261Arg also introduces a new interaction with neighbouring PRPF8 (**Figure 3b; Supplementary Figure 2**). Of the three small substitutions which do not affect H bonding, we discovered that in all cases the variant amino acid was proline, which introduces a new kink in the amino acid chain. Each of these substitutions also resulted in the loss of a polar contact (**Figure 3c,d,e**).

**Figure 3.**
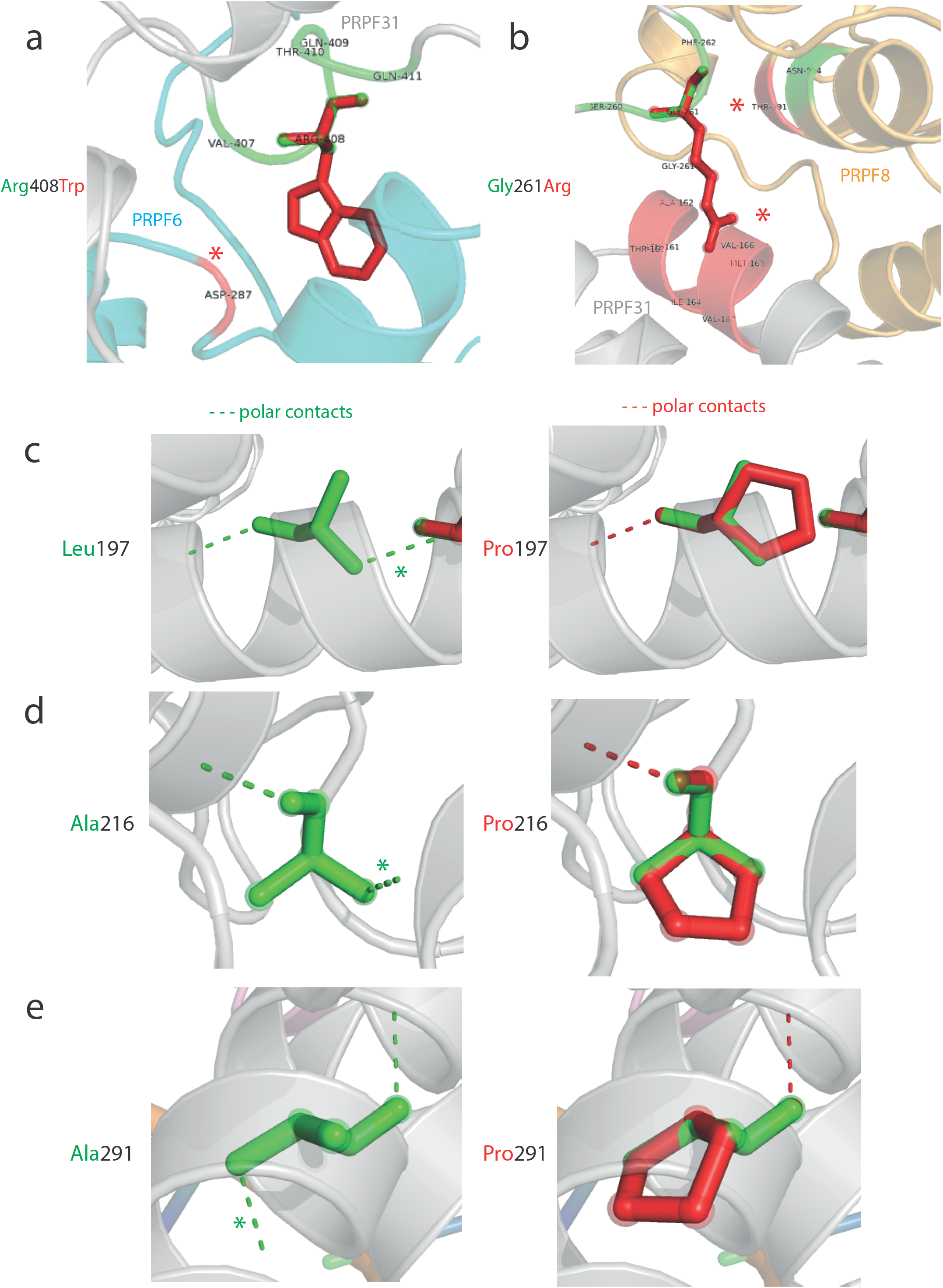
3D cartoon representation of regions of PRPF31 with published missense mutations and their interactions with other molecules within 4Å, and their polar contacts Cartoon representation of alpha helical structure of regions of PRPF31 (grey), with published missense mutations (a) Arg408Trp showing how this affects interaction with PRPF6 (blue) and (b) Gly261Arg showing how this affects interaction with PRPF8 (orange). Red asterisks are used to label where missense mutations introduce new H bonding. Cartoon representation of alpha helical structure of regions of PRPF31 (grey), with published missense mutations (c) Leu197, (d) Ala216Pro and (e) Ala291Pro showing effect of these missense mutation on loss of polar contacts within PRPF31. Wild-type amino acid structure is shown in green, and mutant amino acid structure overlaid in red.

We next mapped the variant found in our patient attending the genetics eye clinic at Bristol Eye Hospital; Ile114Asn (**Figure 4a)**. Ile114Asn is in the coiled-coil domain of the protein, in close proximity to published pathogenic variants Thr138Lys and Ala194Glu (**Figure 4b)**. The substitution introduces new H bonds between this residue and Ala190 of an adjacent α-helix, and predict it to affect protein folding and solubility, and be pathogenic.

**Figure 4.**
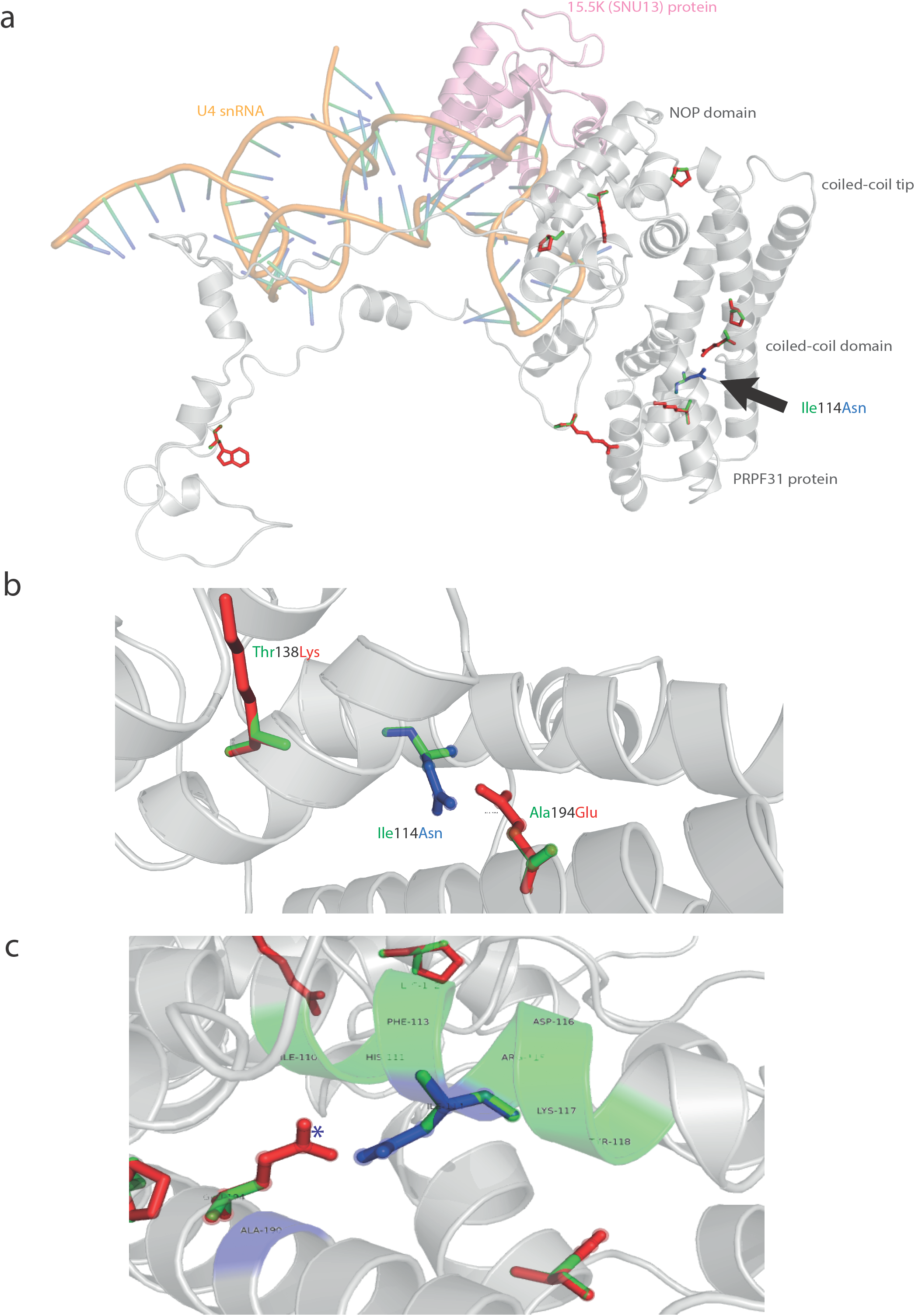
3D cartoon representation of PRPF31 and variant Ile114Asn (a) Cartoon representation of alpha helical structure of PRPF31 (grey) and 15.5K/SNU13 (pink) with U4 snRNA (orange backbone), with published missense mutations mapped onto the physical structure, with wild-type amino acid structure in green, and mutant amino acid structure overlaid in red. Ile114Asn (black arrow) is mapped onto the structure with wild-type amino acid structure in green, and mutant amino acid structure overlaid in blue. (b) Cartoon representation of alpha helical structure of subregion of PRPF31 (grey), with Ile114Asn, showing proximity to Thr138 and Ala194, both of which are published sites of mutation in RP patients (c) Ile114Asn mapped onto the physical structure of PRPF31 with wild-type amino acid structure in green, and mutant amino acid structure overlaid in blue, and interactions within 4Å, predicted to affect H bonding within PRPF31. Green regions of the alpha helix denote normal H bonding by Ile114, blue regions of the alpha helix denote novel H bonds of Asn114. Blue asterisks are used to label where missense mutation introduces new H bonding.

To test the accuracy of our predictions, we took on c.341T>A p.Ile114Asn for further *in vitro* characterisation.

### *In vitro* analysis of c.341T>A p.Ile114Asn variant

To investigate whether c.341T>A p.Ile114Asn caused mislocalisation of the protein, we transfected RPE1 cells with plasmids expressing either wild-type (WT) PRPF31 or PRPF31 341T>A, both tagged with c-myc epitope tag. We used the Lonza nucleofector to ensure high transfection efficiency (∼75%). We assayed the cells after 24, 48 and 72 hours by immunofluorescence confocal microscopy using an anti-cmyc antibody and saw consistent mid- to high-level expression of the WT protein exclusively in the nucleus (**Figure 5a**). We did not observe the same pattern in cells expressing the mutant protein. In these cells, intense c-myc staining was seen in the nuclei of a subset of cells, and no cells showed normal nuclear expression levels (**Figure 5a**). After 72 hours, many cells in the mutant experiment had died, or showed abnormal nuclear morphology (**Figure 5b**). We hypothesised that the mutant protein was aggregating in the nuclei and causing cell death.

**Figure 5.**
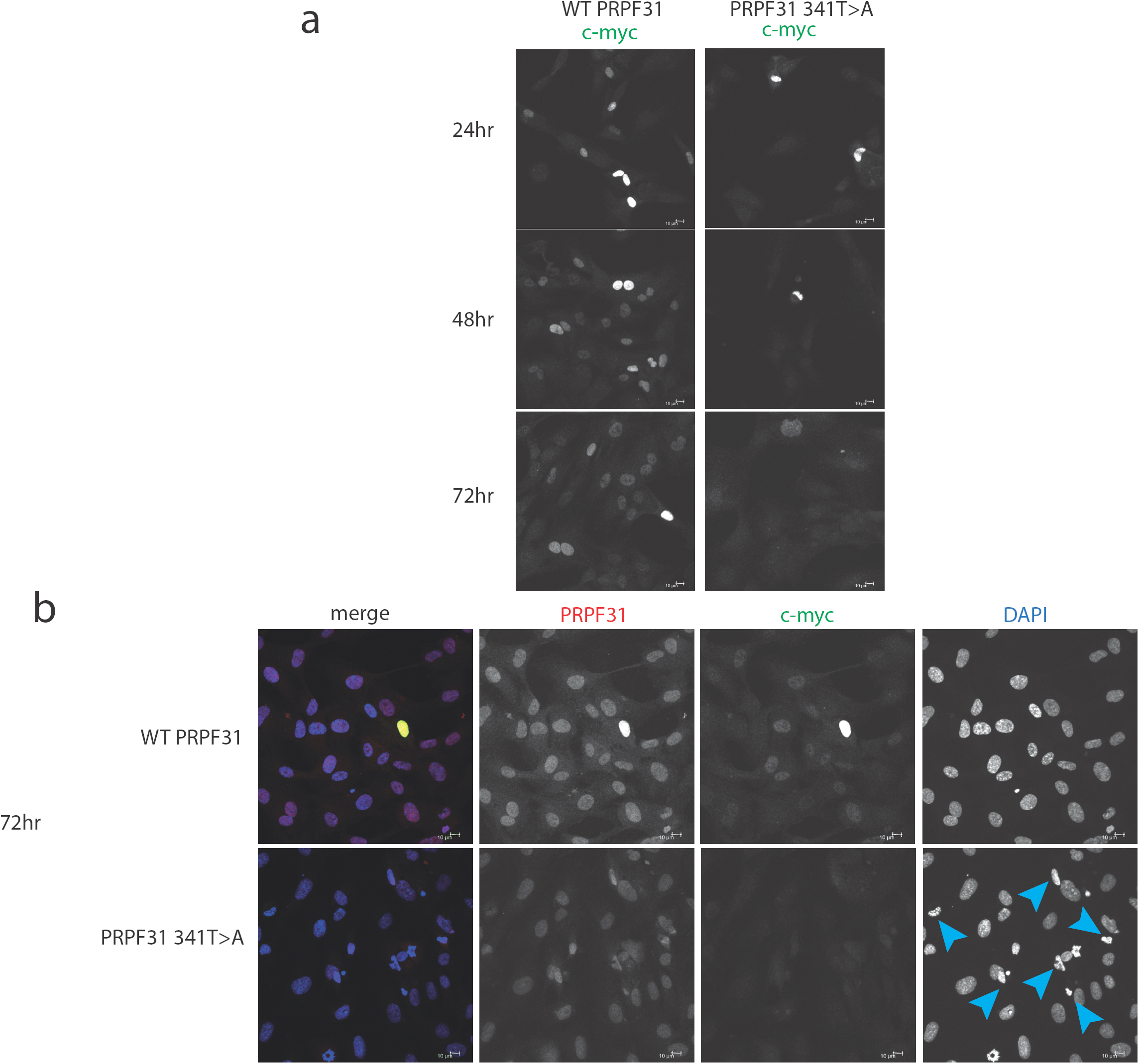
*In vitro* characterisation of *PRPF31* c.341T>A Ile114Asn variant **(a)** Immunofluorescence confocal images of RPE1 cells transfected with c-myc-tagged wild-type or mutant PRPF31, showing expression and localisation of PRPF31-cmyc over 24, 48 and 72 hours. c-myc PRPF31 is evenly distributed throughout the nuclei of cells transfected with WT plasmid at each time point, but is concentrated in the nuclei of a few cells in RPE cells transfected with the mutant plasmid. The number of c-myc positive nuclei is stable in WT cells, but decreases over time in mutant cells. (**b**) At 72 hours, nuclei staining shows many apoptic nuclei (blue arrows) in the cells transfected with mutant PRPF31

In order to investigate whether c.341T>A p.Ile114Asn affected protein stability in a similar way, we transfected HEK293 cells with plasmids expressing either wild-type PRPF31 or PRPF31 341T>A, both tagged with c-myc epitope tag. We treated the transfected cells with cycloheximide protein translation inhibitor over a time course of 6 hours, and assayed protein concentration over this period via western blotting.

Following our usual method for total protein extraction from cells using 1% NP40 detergent, we had difficulty extracting any mutant protein from the transfected cells (**Figure 6a**). This was despite the fact that we could observe protein expression in both cell types via immunofluorescent staining with anti-PRPF31 and anti-cmyc antibodies. We proceeded to repeat the experiment using cell fractionation, to selectively extract protein from the nuclear fraction using 0.1% SDS. This yielded a small amount of mutant protein (**Figure 6b**). Based on our observations, we hypothesised that the mutant protein was in the insoluble nuclear fraction. Once again, we fractionated the cells and lysed the nuclei with 0.1% SDS, but this time we did not remove the insoluble material by centrifugation, instead loading both soluble and insoluble nuclear protein on the gel. This revealed mutant protein, and confirms that the mutant protein is expressed in cells, but is insoluble (**Figure 6c**). No difference in protein stability was observable in the course of cycloheximide treatment (**Figure 6c**). Once we had optimised protein extraction from these cells, we were able to confirm our finding from immunofluorescent imaging that both the WT and mutant protein localised to the nucleus, not the cytoplasm (**Figure 6d**).

**Figure 6.**
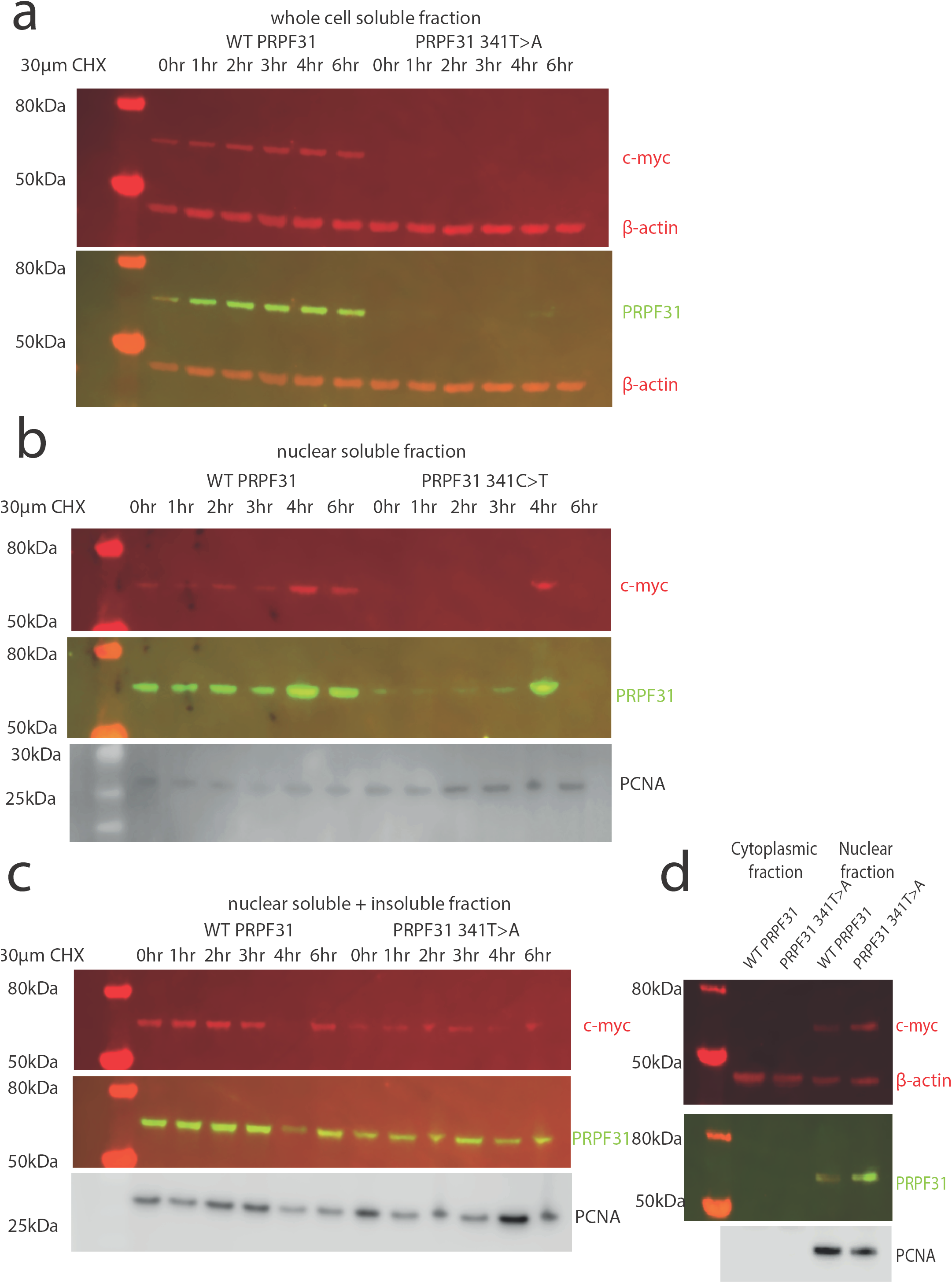
Western blots of protein extracted from HEK293 cells transfected with wild-type or c.341T>A PRPF31 tagged with c-myc **(a)** Cells treated with 30µM cycloheximide (CHX) over 6 hours, and soluble protein extracted from the whole cell showed stable levels of wild-type protein expression across the time course, and complete absence of mutant protein in the soluble whole cell fraction. β-actin is cytoplasmic loading control. **(b)** Cells treated with 30µM cycloheximide (CHX) over 6 hours, and soluble protein extracted from the nucleus showed stable levels of wild-type protein expression across the time course, and extremely low levels of mutant protein in the soluble nuclear fraction, except where some insoluble protein was accidentally loaded (4 hour). β-actin is cytoplasmic loading control. PCNA is nuclear loading control. (**c**) Cells treated with 30µM cycloheximide (CHX) over 6 hours, and both soluble and insoluble protein extracted nucleus showed similar levels of wild-type and mutant protein expression and stability. PCNA is nuclear loading control marker. **(d)** Fractionation shows that both mutant and wild-type PRPF31 are localised to the nucleus. β-actin is cytoplasmic loading control, PCNA is nuclear loading control.

In summary, our findings suggest that c.341T>A p.Ile114Asn variant in *PRPF31* results in protein insolubility, leading to cell death, and is likely the pathogenic cause of RP in this individual. In silico structural analysis of this variant complemented existing techniques for predicting pathoegnecity of this variant.

## Discussion

PRPF31 is a component of the major and minor spliceosome, the huge macromolecular ribonucleoprotein (RNP) complex which catalyses the splicing of pre-messenger RNAs (pre-mRNAs) to remove introns and produce mature mRNAs. More than 90% of human genes undergo alternative splicing (Wang *et al.*, 2008), and splicing is a core function of cells, remarkably well conserved from yeast to man. The spliceosome is composed of at least 43 different proteins, and 5 small nuclear RNAs (snRNAs), U1-U5 (Will and Luhrmann, 2011).

PRPF31 is essential for the assembly of the U4/U6.U5 tri-snRNP complex (Makarova *et al.*, 2002), which, when combined with U1 and U2, forms the ‘B complex’. After large rearrangements, the activated B complex is able to initiate the first step of splicing. In the absence of PRPF31, U4/U6 di-snRNP accumulates in the splicing-rich Cajal bodies in the nucleus, preventing formation of the tri-snRNP, and subsequently efficient splicing (Schaffert *et al.*, 2004).

PRPF31 performs its function through several important protein domains; the flexible loop, Nop domain, coiled-coil domain and tip. The flexible loop (residues 256 – 265) protects the exposed C4’ atoms of residues 37 and 38 from attack by free radicals, to protect the RNA without directly contacting it (Liu *et al.*, 2007).

The Nop domain is a conserved RNP-binding domain, with regions for binding protein and RNA. Although the sequence conservation of the Nop domain is relaxed in PRPF31, its specificity for binding U4 or U4atac and 15.5K protein is high (Liu *et al.*, 2007).

Nonsense, missense and indel mutations in PRPF31 are associated with autosomal dominant RP with incomplete penetrance. Whilst the pathogenicity nonsense and indels is easy to ascribe, the pathogenicity of missense mutations in this protein are difficult to predict. Alongside use of pathogenicity predictors based on 2D structure and conservation, such as PolyPhen, SIFT and CADD, *in silico* analysis of the 3D crystal structure of PRPF31 can provide more accurate estimates of the pathogenic potential of missense mutations. However, this still only provides information about the effect of mutations on the protein in isolation. The elucidation of the structure of the entire intact spliceosomal complex provides new exciting opportunities for more accurate *in silico* modelling of the effect of missense mutations on PRPF31 and its protein-protein interactions (Bertram *et al.*, 2017). This structural analysis can predict effects on interactions with other proteins, as well as intramolecular disturbances caused by missense mutations. For example, one published variant in PRPF31 is in the C-terminal domain (Arg408Trp), outside of the functionally important Nop and coiled-coil domains and in silico analysis predicts that this missense mutation does not affect H bonding within PRPF31. Based on this analysis, it may be predicted that this change is unlikely to be pathogenic, but 3D analysis of the intact spliceosome predicts that this changes affects binding of PRPF31 to PRPF6 (**Figure 3a**). We would predict that this change is pathogenic, and that missense variants outside the Nop and coiled-coil domains should not be dismissed as benign.

Using this 3D protein complex analysis, we predict a novel variant, Ile114Asn, found in a patient attending the genetics eye clinic at Bristol Eye Hospital, to be affecting H bonding within PRPF31 and predict that this will affect protein folding and solubility (**Figure 4a-c**). Our *in vitro* studies confirm this (**Figures 5, 6**). Protein with this variant is found in the insoluble nuclear fraction, and this leads to cell death (**Figures 5, 6**).

In summary, we show that *in silico* modelling of the effect of missense variants on the 3D structure of the spliceosome contributes useful additional data to predictions of pathogenicity of novel variants. which are likely to affect protein folding and solubility. In the novel variant studied here, the predictions from this *in silico* structural analysis were confirmed using *in vitro* testing. It is important to note that the spliceosome is a highly dynamic structure, and our 3D structural complex analysis only studies PRPF31 in one specific conformation, in the spliceosome primed for splicing (Bertram *et al.*, 2017). For truly accurate predictions of pathogenicity, the 3D structure of the spliceosome at different stages of activity will need to be studied, preferably using Molecular dynamic simulation (MDS) with a package such as GROMACS (Berendsen *et al.*, 1995) to provide deepest insights into effects of missense mutations. The publication of more cryo-EM resolved complexes relevant to development of ciliopathies, such as the intraflagellar transport (IFT) complexes (Jordan *et al.*, 2018) will further enhance our understanding of such conditions, and allow more accurate computational prediction of pathogenicity of variants.

Our data from this novel variant supports previous suggestions that haploinsufficiency is the common disease mechanism in RP11 rather than any dominant negative effects of missense variants (Wilkie *et al.*, 2008; Abu-Safieh *et al.*, 2006; Sullivan *et al.*, 2006). We find that this missense variant affects protein solublilty, effectively removing one copy of the protein from cells.

Considerable further work is required to elucidate why haploinsufficiency of PRPF31 causes retinal cells to degenerate, whether specific or global pre-mRNA splicing is affected, and why other tissues outside the retina are not affected by loss of protein.

## Supporting information

**Supplementary Figure 1. 3D cartoon representation of PRPF31, including published missense mutations, in complex with U4 snRNA, 15.5K and PRPF6 (a)** Cartoon representation of alpha helical structure of PRPF31 (grey) and 15.5K/SNU13 (pink) with U4 snRNA (dark orange backbone), and PRPF6 (blue) with published missense mutations mapped onto the physical structure, with wild-type amino acid structure in green, and mutant amino acid structure overlaid in red. This shows that only Arg408Trp is in interacting proximity with PRPF6. **(b)** An alternative view of the same complex, highlighting that variants in the NOP domain (black arrow) and coiled-coil domain do not appear to interact with PRPF6 in this conformation

**Supplementary Figure 2. 3D cartoon representation of PRPF31, including published missense mutations, in complex with U4 snRNA, 15.5K, PRPF6 and PRPF8 (a)** Cartoon representation of alpha helical structure of PRPF31 (grey) and 15.5K/SNU13 (pink) with U4 snRNA (dark orange backbone), PRPF6 (blue), and PRPF8 (light orange) with published missense mutations mapped onto the physical structure, with wild-type amino acid structure in green, and mutant amino acid structure overlaid in red. This shows that only Gly261Arg is in interacting proximity with PRPF8. **(b)** An alternative view of the same complex, highlighting that only this Gly261Arg variant (black arrow) appears to interact with PRPF8 in this conformation

## Author Contributions

GW and AC conceived of and designed the study.

NM, SH, NJ and AC examined the patient, coordinated genetic testing and analysed patient genetic data.

GW and LN carried out in silico and in vitro experiments. GW, LN, NM and AC prepared figures.

GW and AC wrote the paper.

SH reviewed the paper.

