## Supplementary figures and images for "A combined *in silico*, *in vitro* and clinical approach to characterise novel pathogenic missense variants in PRPF31 in retinitis pigmentosa"

### Supplementary file 1

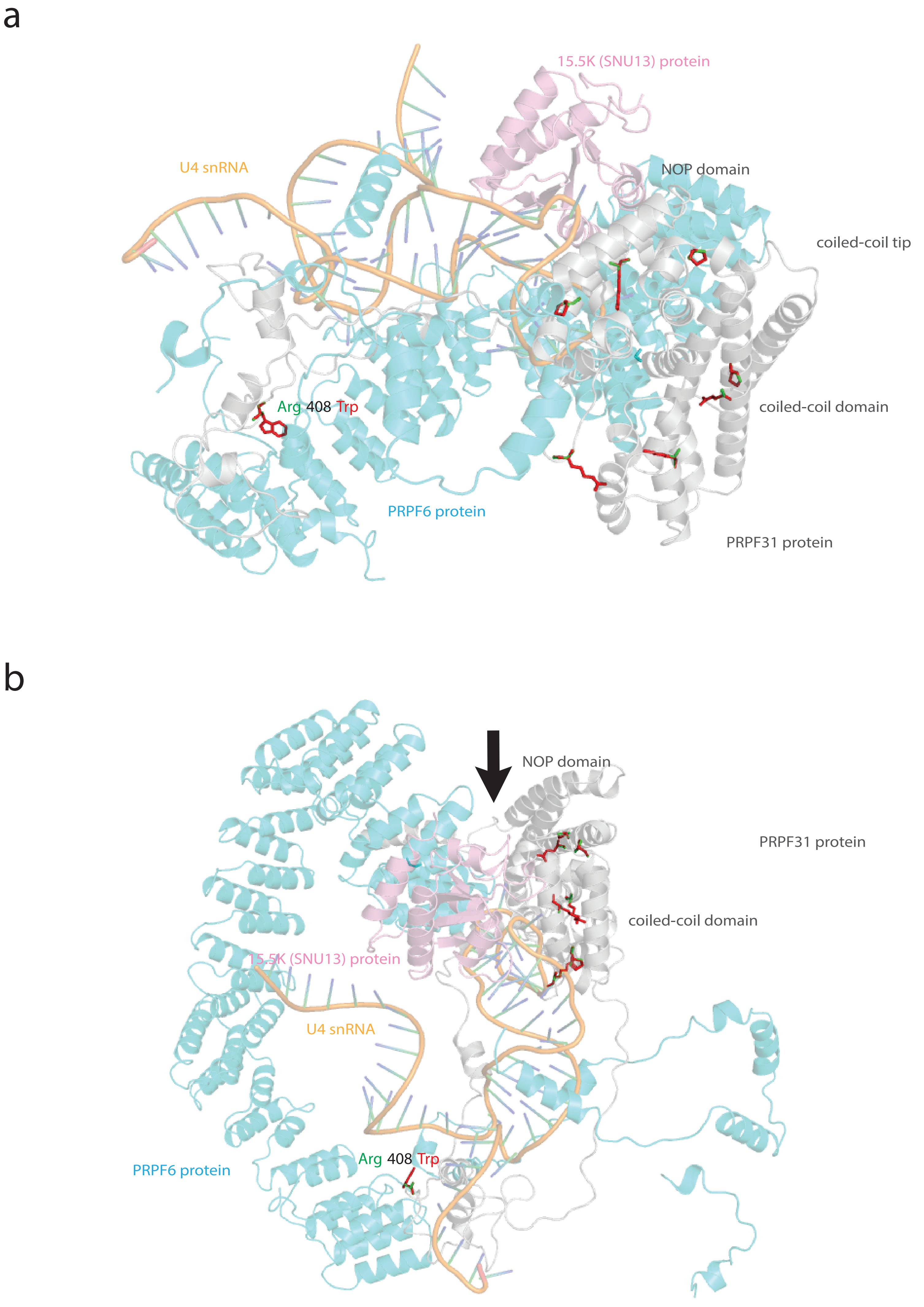

### Supplementary file 2

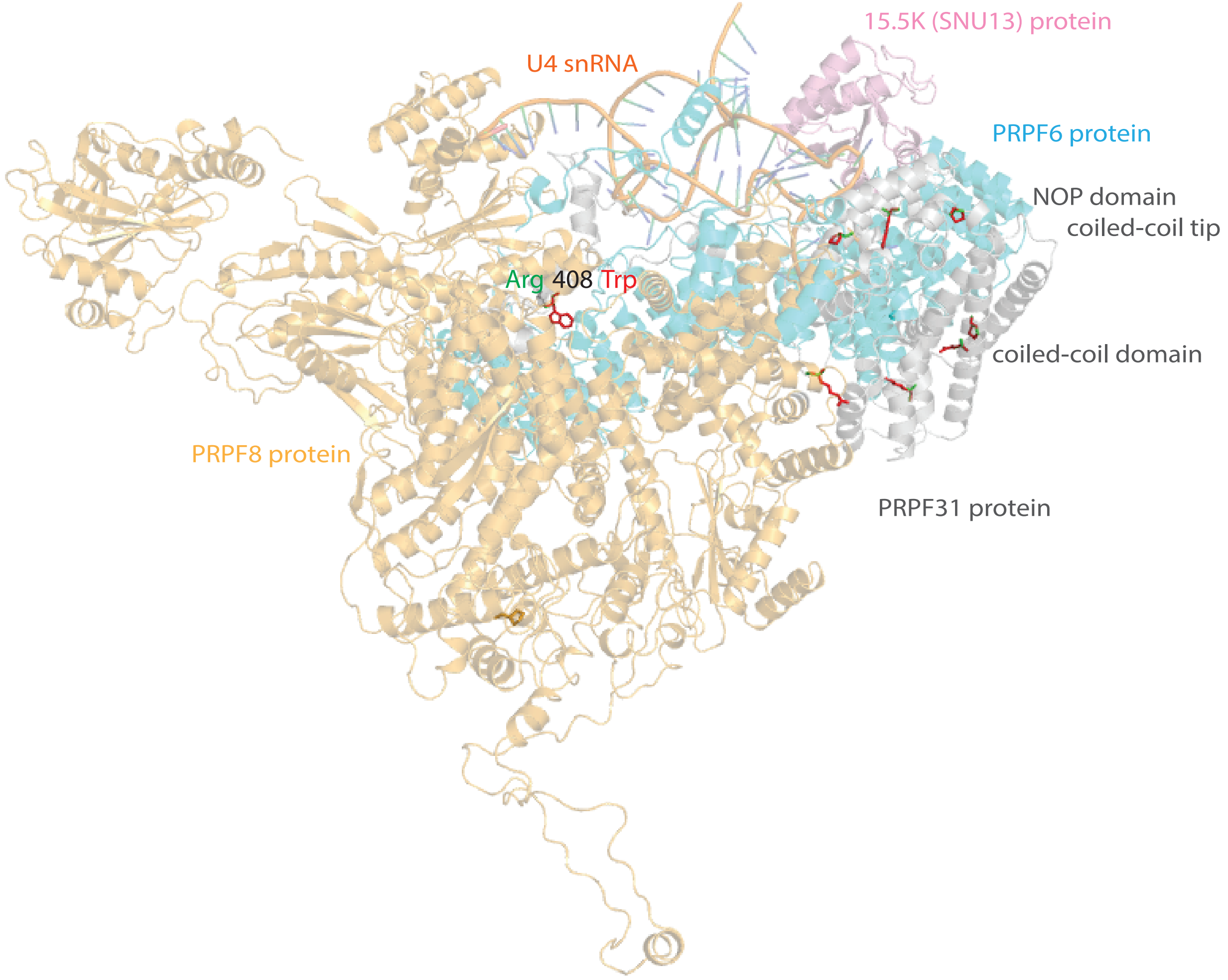

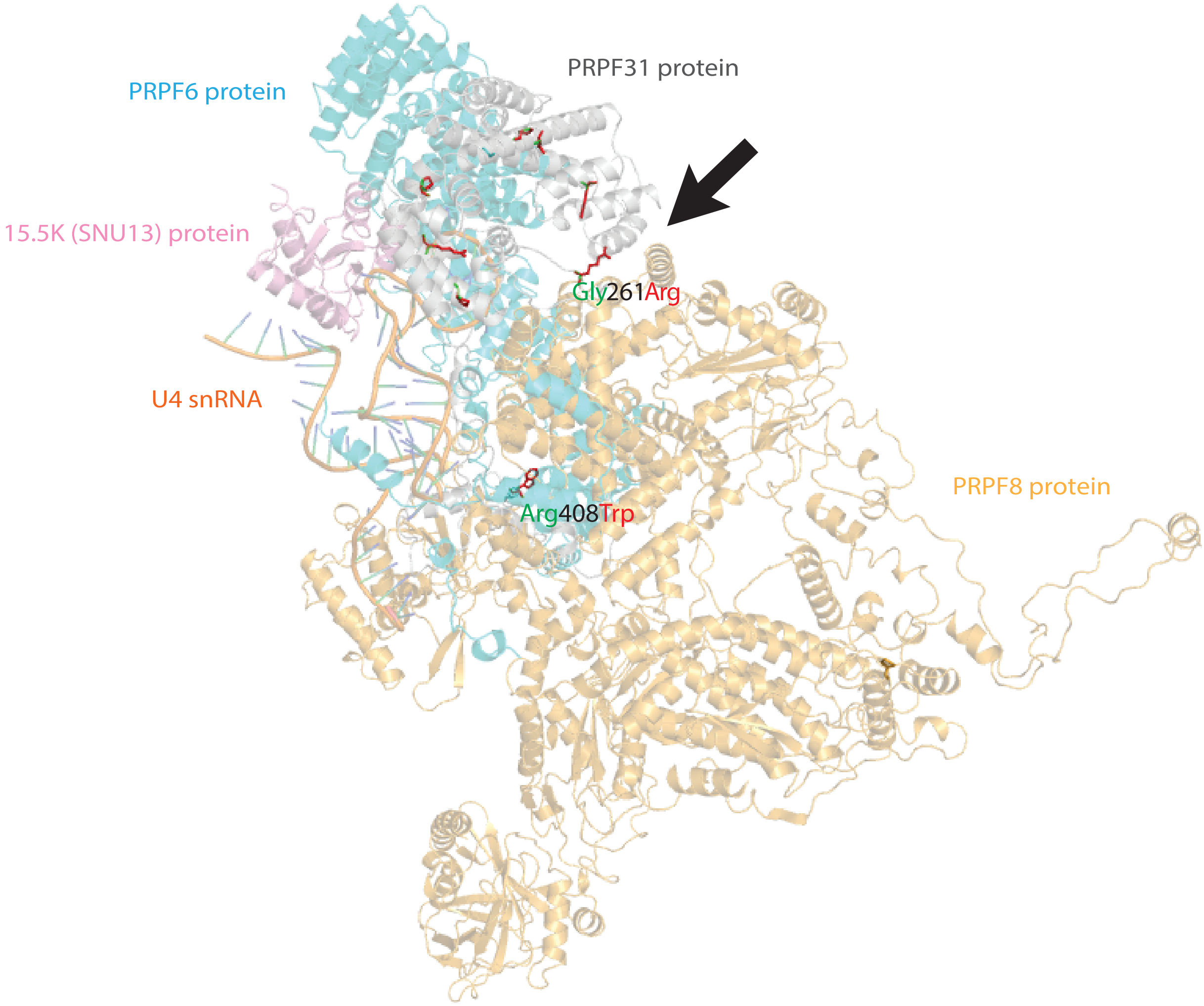
